# Multiplexing of temporal and spatial information in the lateral entorhinal cortex

**DOI:** 10.1101/2024.01.31.578307

**Authors:** Cheng Wang, Heekyung Lee, Geeta Rao, James J. Knierim

## Abstract

Episodic memory involves the processing of spatial and temporal aspects of personal experiences. The lateral entorhinal cortex (LEC) plays an essential role in subserving memory. However, the specific mechanism by which LEC integrates spatial and temporal information remains elusive. Here, we recorded LEC neurons while rats performed foraging and shuttling behaviors on one-dimensional, linear or circular tracks. Unlike open-field foraging tasks, many LEC cells displayed spatial firing fields in these tasks and demonstrated selectivity for traveling directions. Furthermore, some LEC neurons displayed changes in the firing rates of their spatial rate maps during a session, a phenomenon referred to as rate remapping. Importantly, this temporal modulation was consistent across sessions, even when the spatial environment was altered. Notably, the strength of temporal modulation was found to be greater in LEC compared to other brain regions, such as the medial entorhinal cortex (MEC), CA1, and CA3. Thus, the spatial rate mapping observed in LEC neurons may serve as a coding mechanism for temporal context, allowing for flexible multiplexing of spatial and temporal information.

## Introduction

Our memory of everyday experiences involves proper representation of spatial and temporal context. The hippocampal memory system plays an essential role in episodic memory. Indeed, diverse mechanisms for representing space and time have been revealed in the hippocampus and its neighboring regions ^1–4^. However, the exact neural mechanisms responsible for representing space and time upstream of the hippocampus remain unknown.

The lateral entorhinal cortex (LEC) and medial entorhinal cortex (MEC) constitute the major cortical inputs to the hippocampus, and they exhibit important anatomical and functional specializations ^5,6^. MEC contains functional cell types such as grid cells, boundary/border cells, head direction cells, non-grid spatial cells, and speed cells ^7–11^. The presence of these cells strongly supports the hypothesized role of the MEC in constructing a universal, allocentric map through the computation of path integration ^12–14^. In contrast, LEC neurons display comparatively little allocentric spatial information ^15,16^. LEC neurons receive rich multisensory information about the external world and can respond to olfactory, visual, and auditory stimuli ^17–19^. LEC cells encode spatial information related to the objects or items in the environment ^20–23^, and they appear to primarily represent this information in an egocentric coordinate frame ^22,23^. Additionally, LEC is involved in encoding temporal information across time scales spanning seconds to hours ^24^. This finding is corroborated by the evidence of a positive correlation between blood-oxygen-level-dependent activity in LEC and the precise temporal position judgment of movie-frames within a movie by human subjects ^25^. Thus, LEC encodes forms of both spatial and temporal information, but how LEC neurons integrate these types of information remains largely unexplored.

To address these questions, we characterized the neural activity of LEC while rats ran on 1-D linear and circular tracks. We found that many LEC neurons showed robust spatial selectivity in these tasks, showing sensitivity to both the direction of travel and manipulation of the spatial environment. In addition, the spatial rate maps demonstrated temporal modulation across laps within a session, suggesting the representation of temporal information through spatial rate remapping. The results reveal a flexible, spatially and temporally multiplexed code in LEC.

## Results

### Behavioral Setup

We used three types of one-dimensional tasks: the double rotation task, the circular track task, and the linear track task. In the double rotation task ^26^, rats moved clockwise on a circular track for irregularly placed food reward; local cues on the track and global cues on a curtain in the periphery of the room were put into conflict during Mismatch sessions (Fig. 1a). In the circular track and linear track tasks (Fig. 1b and 1c), food wells were fixed at the end of the journey, and the rats were required to traverse the same locations in two different directions. For the circular track task, a dark session in which lights were turned off was placed between two standard (Light) sessions. These task structures enabled us to investigate the representation of spatial and temporal information in LEC. A subset of the data was reported in prior studies ^24,27^ to address questions that were different from the present report.

**Figure 1.**
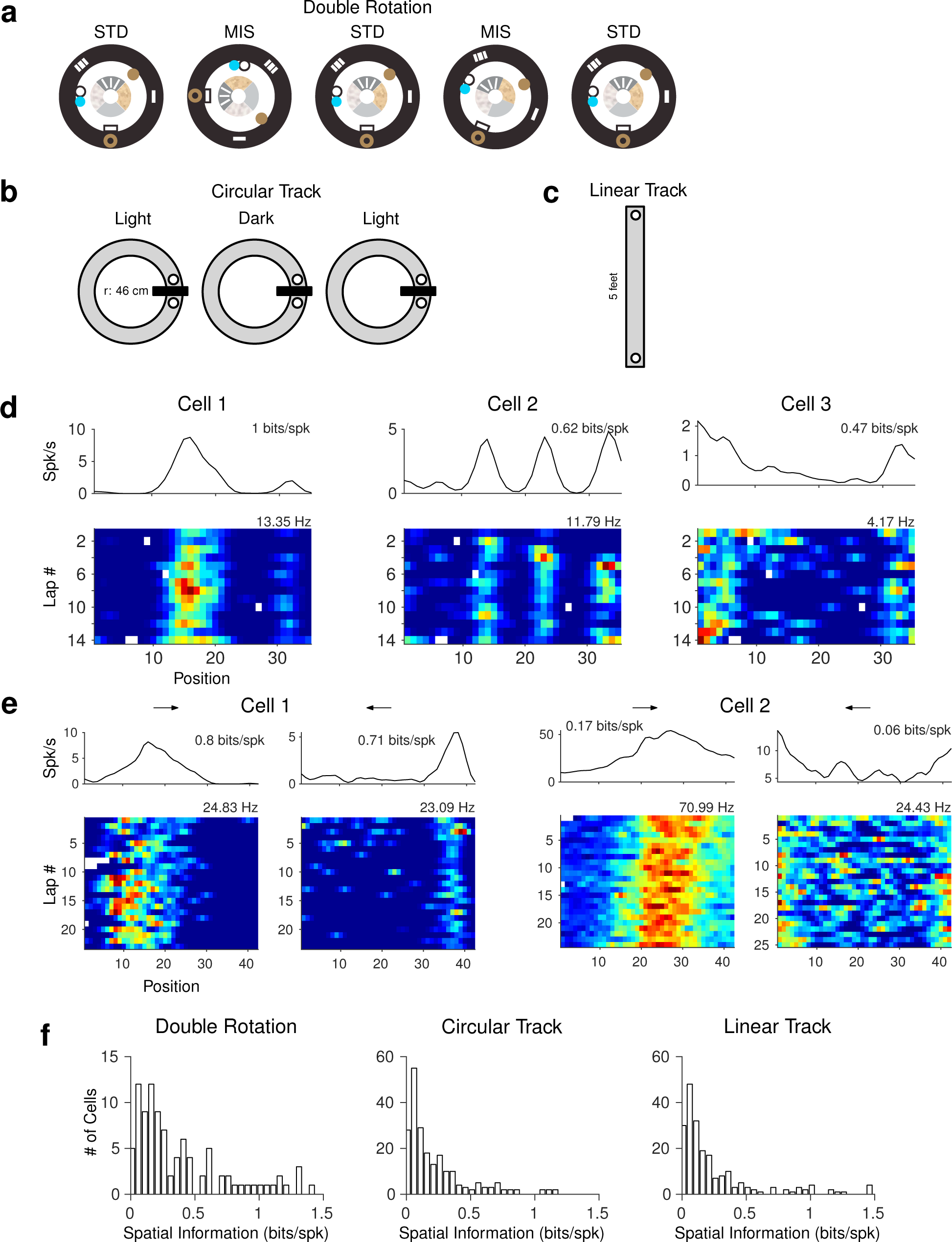
Spatial firing of LEC neurons in one-dimensional apparatus. (**a**) Schematics of the double rotation task. Three standard (STD) sessions were interleaved with two mismatch (MIS) sessions. In the MIS sessions, local cues on the track were rotated counterclockwise and the global cues along the curtain at the periphery of the room (the black outer ring) were rotated clockwise. The subjects foraged for food scattered at arbitrary locations on the track (∼2 rewards/lap) while moving clockwise. (**b**) The circular track task. Two sessions recorded in the light condition were separated by a session in the dark. (**c**) Linear track. For both the circular track and linear track, the animals shuttled back and forth to retrieve food pellets in the food wells placed at the ends of the journeys. (**d**) Spatial firing patterns of three example LEC neurons in the double rotation task. For each cell, the spatial rate map of the session is shown at the top (the number on top indicates the spatial information score) and the lap-wise spatial rate map is shown at the bottom (the number on top shows the peak firing rate of the map). (**e**) The session-wise spatial rate maps and lap-wise spatial rate maps of two LEC cells in the circular track task. The rate maps of the two movement directions are separately shown for each cell. (**f**) Histograms of the distribution of spatial information scores for all LEC neurons in the three behavioral tasks.

### LEC neurons showed spatial selectivity in one-dimensional tasks

We recorded a total of 368 LEC neurons from 8 male Long-Evans rats (Fig. S1) in the three tasks (119 neurons from 3 rats in the double rotation, 239 neurons from 5 rats in the circular track, and 231 neurons from 5 rats in the linear track task; some neurons were recorded over multiple tasks). We excluded cells that fired < 20 spikes within a session (defined as minimally active cells) or whose firing rate was > 10 Hz (putative interneurons ^15^). To characterize the spatial firing patterns of these neurons in one-dimensional tasks, we first linearized the position in the circular track task and double rotation task. There were frequent off-track head scanning events in the double rotation task. We detected and removed position frames during these scanning events (Fig. S2a), as these events were typically spatially inhomogeneous and behaviorally distinct from the dominant forward movement behavior in these tasks ^28^. Surprisingly, unlike previous studies of LEC in empty, 2-D arenas that showed minimal amounts of allocentric spatial representation ^15,16^, some LEC cells in all three tasks showed clear spatial firing fields in allocentric rate maps (Fig. 1d to 1f; Fig. S2b to 2d). For some neurons, the spatial selectivity and reliability were similar to those of hippocampal neurons. Overall, the median spatial information score in the double rotation task was 0.27 (IQR = 0.14 – 0.60); in the circular track task was 0.141 (IQR = 0.077 – 0.336); and in the linear track task was 0.140 (IQR = 0.073 – 0.341).

We next investigated how different manipulations of the spatial context (cue-mismatch sessions in the double rotation task and light-dark sessions in the circular track task) influenced the spatial firing of LEC cells in these tasks (Fig. 2a and 2b). First, the spatial selectivity of neurons was preserved despite the large (135° and 180°) mismatch of local and global cues in the double rotation task (median information score: STD = 0.22, IQR = 0.13 – 0.44; MIS = 0.27, IQR = 0.15 – 0.57; Wilcoxon rank-sum test, *Z =* −0.27, *p =* 0.79) or the absence of visual inputs in the circular track task (median information score: STD = 0.21, IQR = 0.09 – 0.39; Dark = 0.22, IQR = 0.10 – 0.49; Wilcoxon rank-sum test, *Z =* −1.23, *p =* 0.46). However, the Pearson correlation coefficients between rate maps in the standard session and the manipulated session were significantly smaller than those between the two standard sessions (Fig. 2c and 2f; double rotation task: median difference = 0.36, IQR = 0.06 − 0.72; circular track task: median difference = 0.11, IQR = −0.029 − 0.31; Wilcoxon signed-rank test, double rotation task: *p* < 0.001, *Z* = 5.41; circular track task: *p* < 0.001, *Z* = 5.16). To further visualize the results, population correlation matrices were created as previously described ^27,29,30^, and they demonstrated that the spatial firing patterns were largely preserved across manipulations (Fig. 2d and 2e, Fig. 2g). In the double rotation task, the band of high correlation shifted downward from the main diagonal in the STD 1 vs. MIS matrices, indicating that the spatial firing fields were dominated by the local cues on the track (as previously demonstrated in this data set ^27^, although the strength of the correlation band in the 180° mismatch session is weaker than the 135° mismatch session). The reduced correlations of individual place fields in Fig. 2c result partially from the rotation of the firing fields in the mismatch session. In the circular track, the correlation matrices were very similar between the light and dark sessions, although the correlation bands were more diffuse in the dark. Overall, these results demonstrate that the spatial representations of LEC neurons were sensitive to changes in the environmental context.

**Figure 2.**
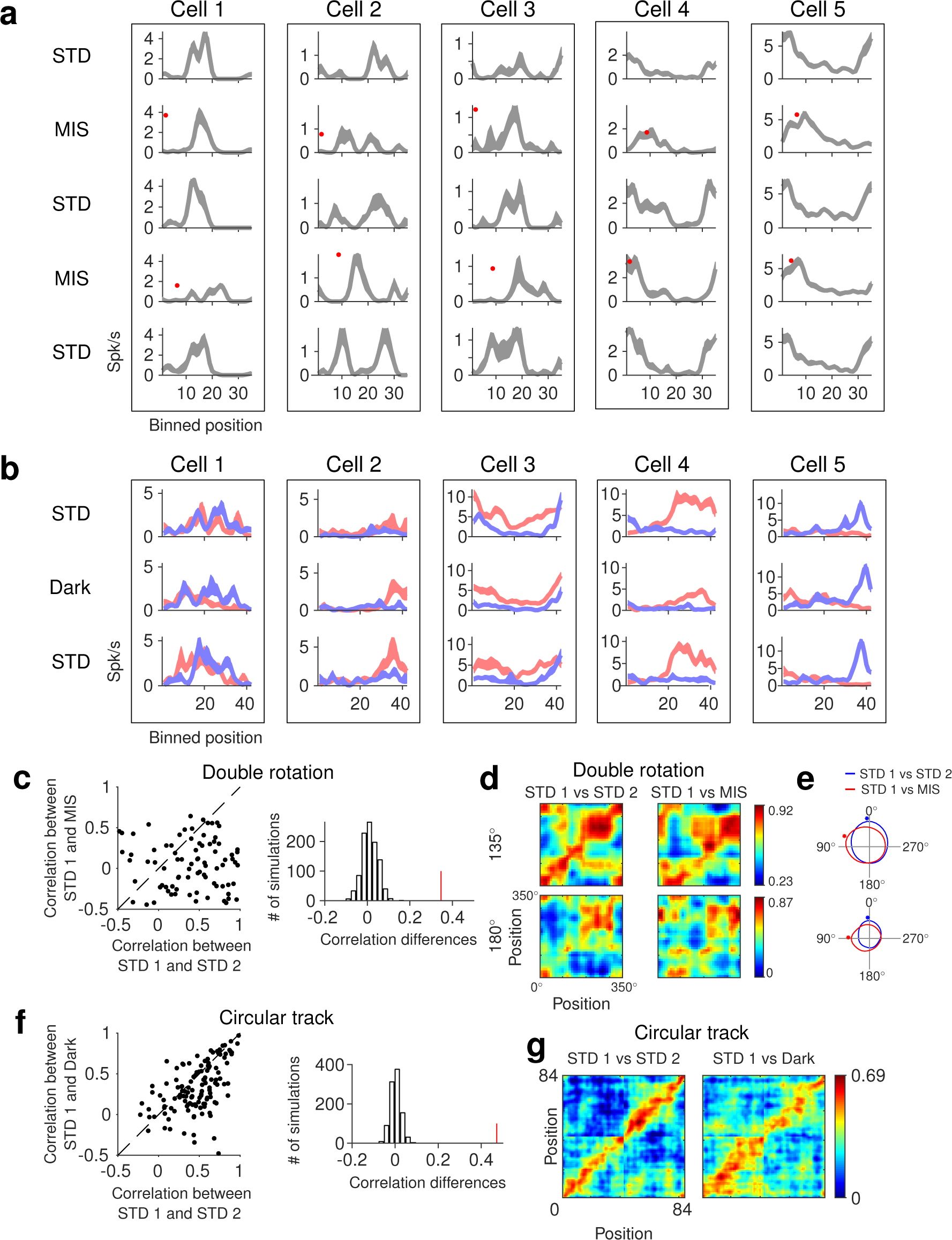
The influence of environmental manipulations on the spatial firing of LEC neurons. (**a**) Spatial rate maps for five example LEC neurons in the 5 sessions of the double rotation task. For each rate map, the shaded region indicates ±1 S.E.M. around the mean firing rate. The red dots denote the positions (under the camera) of the 0° locations on the STD tracks after cue manipulations in the MIS sessions. Qualitatively, cells show similar magnitudes of spatial tuning in the standard and mismatch sessions, although the firing rate peaks shift between sessions, often under the control of the local cues on the track, e.g., cell 4 and 5 (Neunuebel et al., 2013). (**b**) Same as panel a but for the spatial firing of five LEC cells in each of the two directions (blue: CW and red: CCW) in the circular track task. Spatial firing fields have different profiles for CW vs. CCW directions (see Figure 3), but the fields are qualitatively similar across light and dark conditions. (**c**) Left, scatter plot of the correlation between two STD sessions and between STD and MIS sessions. Right, the observed difference in correlations (the red line) between the two groups (STD1 vs. STD2 and STD1 vs. MIS) is outside the range of the null distribution of correlation differences derived from a permutation analysis. (**d**) Population correlation matrices between two sessions in the double rotation task for 135° (top) and 180° (bottom) mismatch sessions. Left: STD1 vs. STD2, right: STD1 vs. MIS. Although a similar analysis was performed in a previous study using the same dataset from the double rotation task, this analysis was repeated as we changed the method for computing spatial rate maps (completely removing head scanning events and changing the spatial bin size, see Methods). The downward shift of the correlation band in the STD1 vs. MIS matrix demonstrates that the spatial firing of LEC cells tended to follow the rotation of the local cues (Neunuebel et al., 2013). (**e**) Polar plots created by averaging the bins along the diagonals of the correlation matrix. Top and bottom, polar plots for 135° and 180° manipulations, respectively. The asterisks indicate the angles with the largest correlation. The peak angles of the STD 1 vs. MIS polar plots are approximately 67.5° and 90° CCW for the 135° and 180° manipulations, respectively, indicating that the LEC representation was predominantly controlled by the local cues (Neunuebel et al., 2013). (**f**) and (**g**), same as panels c and d for STD and Dark sessions in the circular track task. The spatial rate maps of the two movement directions were concatenated.

### The spatial firing of LEC neurons was sensitive to movement direction

Hippocampal and medial entorhinal cortex neurons show direction selectivity when rats perform shuttling tasks on one-dimensional tracks ^31–36^. We found that the spatial firing of LEC neurons also distinguished traveling directions in the circular track and the linear track tasks (Fig. 3a and Fig. S3). For some neurons, the overall firing was greater for one direction than the other; for other neurons, the cell preferred different directions at different locations on the track. A permutation method was adapted from Fujisawa et al. ^37^ to identify the cells with significant direction selectivity (see Methods). Approximately 40% of LEC neurons (circular track task: 97/213, linear track task: 90/211) had a significant preference for travel direction in at least one location. The distribution of directionally selective locations is significantly different from a uniform distribution, as the locations with directional selectivity were biased toward the ends of the track (Fig. 3b, Rayleigh test, *p* < 0.001 for both standard and dark circular track data; Fig. 3c, Monte Carlo test, *p* < 0.001 for linear track data).

**Figure 3.**
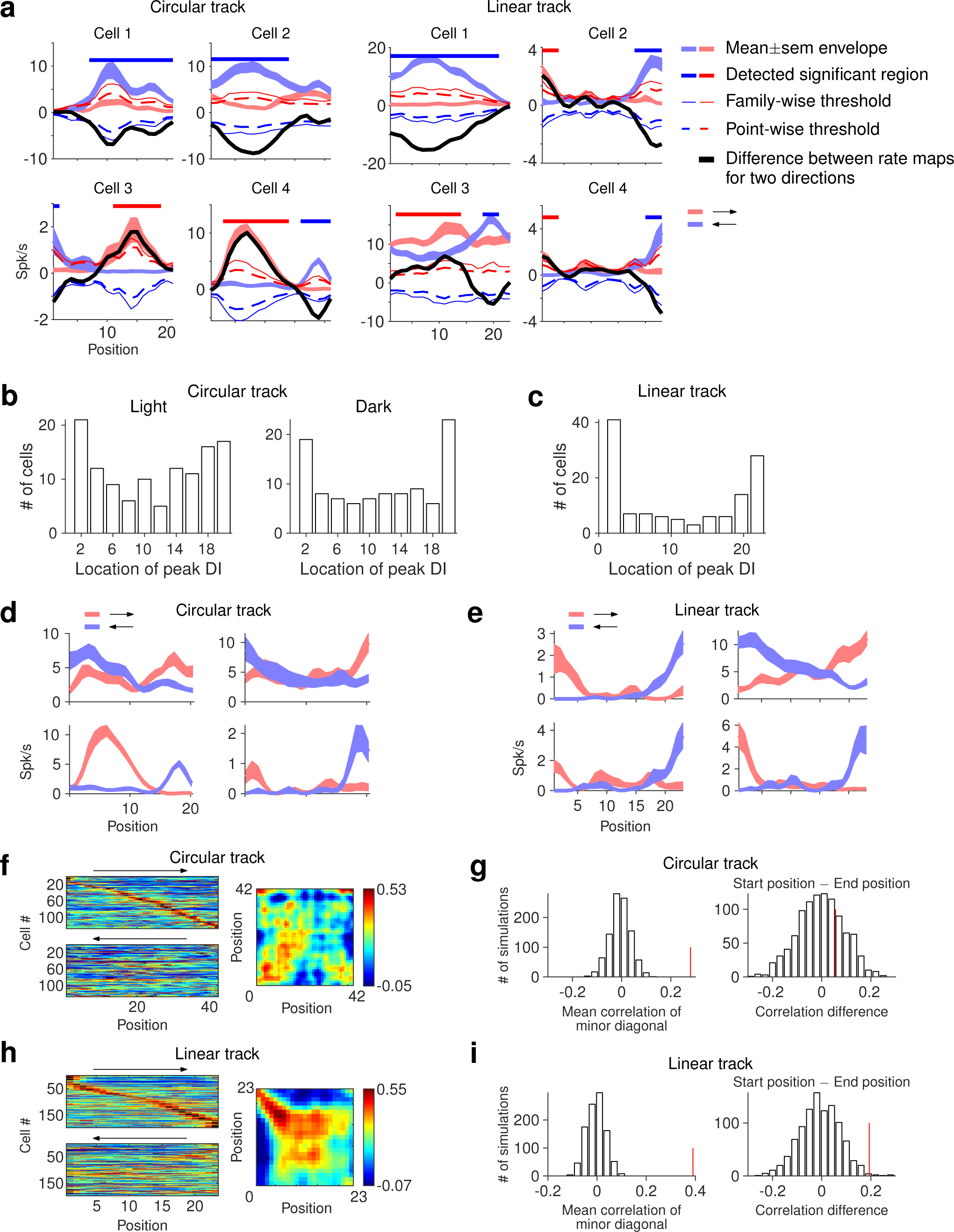
Directional preferences of LEC cells in 1-dimensional shuttling tasks. (**a**) Left, four example LEC neurons showing directionally selective firing on the circular track. Shaded red and blue colors denote the mean ± SEM for counterclockwise and clockwise movement, respectively. The black line denotes the difference between the firing rates for the two directions. Colored lines and dotted lines are thresholds for detecting pointwise and family-wise significant regions, respectively. Bold color lines at the top denote regions of significant direction selectivity. For cell 1 in the circular track task, the cell had significantly higher firing when moving in the blue direction for most of the track. The black line crosses the lower bound of the family-wise threshold (blue line), indicating the cell has significant direction selectivity at this location, and the spatial extent of direction selectivity is determined by the region in which the black line is below the lower bound of the point-wise threshold (blue dashed line). Cell 3 and cell 4 in the circular track task both prefer different directions at different locations on the track. Right, four neurons with direction selectivity in linear track paradigm. Same conventions as for the circular track (left). (**b**) Distribution of locations with peak direction selectivity index (see Methods) for each cell on the circular track. (**c**) Same as Figure 3b for the linear track. (**d**) Four example LEC neurons with strong distance coding in the circular track task. All four cells fired comparably depending on the distance from the start of the journey. (**e**) Same as panel d for four cells in the linear track task. (**f**) Left: sorted spatial rate maps for the two directions in the circular track task. Right: population correlation matrix between the two directions. A high correlation in the main diagonal (bottom left to top right) indicates consistent firing at the same location for the two directions, whereas a strong signal in the minor diagonal (top left to bottom right) indicates consistent firing at the same journey distance from the track ends. (**g**) Left, the observed mean correlation of bins along the minor diagonal of the correlation matrix (i.e., from top left to lower right showing the correlation between the same distances from the start position in both directions) is shown as the red line and the null distribution is simulated with a permutation analysis. Right, the observed correlation difference (the red line) and the null distribution comparing the distance coding from the starting position (the first 4 bins) and the end position (the last 4 bins). (**h**) and (**i**) Same as panels f and g for the linear track task.

In some cases in which a cell preferred different directions at different locations on the track, the locations were symmetrically organized (Fig. 3d and 3e; Fig. 3f and 3h). In other words, the cells appeared to be sensitive to the distance traveled along the track in each direction. Fig. 3f and 3h show the rate maps of all cells on the circular and linear tracks, respectively, in each direction, ordered by the peak firing rate in the first direction. The lack of a strong diagonal in the ordered rate maps of the opposite direction demonstrates the prevalence of directional firing. To the right of the ordered rate maps are correlation matrices of the ordered rate maps for each direction. For the circular track (Fig. 3f), the correlation matrix shows an X-shaped pattern, indicating that some cells fired at the same locations in both directions (the main diagonal), whereas other cells showed distance coding (the minor diagonal). The average correlation along the minor diagonal was significantly larger than a null distribution (Fig. 3g, left, *p* < 0.001). Similar results were observed on the linear track (Fig. 3h; Fig. 3i, left, *p* < 0.001), although there was a strong asymmetry between the strength of the correlations on the upper left and the lower right quadrants of the correlation matrix. This asymmetry indicates that the distance coding was significantly stronger at the start of the journey than at the end, a result not observed in the circular track task (circular track: Fig. 3g right, *p* = 0.29; linear track: Fig. 3i right, *p* = 0.007). However, a significant difference in the differential distance tuning for the start vs. the end of the journey was not observed between the linear track and the circular track (*p* = 0.24, permutation test, not shown). The lack of a statistical difference between linear and circular track data made the apparent difference in the correlation matrices ambiguous.

### Representation of trial progression by LEC neurons

Because the entorhinal cortex has been implicated in the processing of temporal information ^24,25,38–41^, we explored whether LEC cells encode information about the temporal progression of trials. When lap-wise spatial rate maps were created for LEC, the spatial firing rates for different laps showed clear modulation across the session (Fig. 4 and Fig. S4). We used singular value decomposition (SVD) to decompose the lap-wise spatial rate maps into spatial and temporal firing profiles, which we termed spatial modulation fields and temporal modulation fields, respectively (Fig. 5a, Fig. S5; see Methods). Fig. 5b and 5c demonstrate the reproducibility across sessions of temporal modulation fields for 12 neurons with highly reproducible temporal firing profiles in the double rotation and circular track tasks (see Fig. S4 for examples of other cells with less reproducible firing). We quantified this reproducibility by calculating correlation coefficients of temporal modulation fields across sessions for each neuron. The mean correlation coefficients of all neurons were significantly greater than those from null distributions obtained by permuting cell labels in the second session (the double rotation task: Fig. 5d, STD 1 vs. STD 2, *p* = 0.022, STD 1 vs. STD 3, *p* = 0.018; the circular track task: Fig. 5e, direction 1, *p* = 0.011, direction 2, *p* < 0.001). Some LEC cells showed consistent temporal modulation in the absence of strong spatial selectivity (Fig. S4), which suggests that a temporal signal could be present independent of spatial selectivity^24^. Altogether, our results demonstrate that LEC might represent temporal information about trial identity through spatial rate remapping.

**Figure 4.**
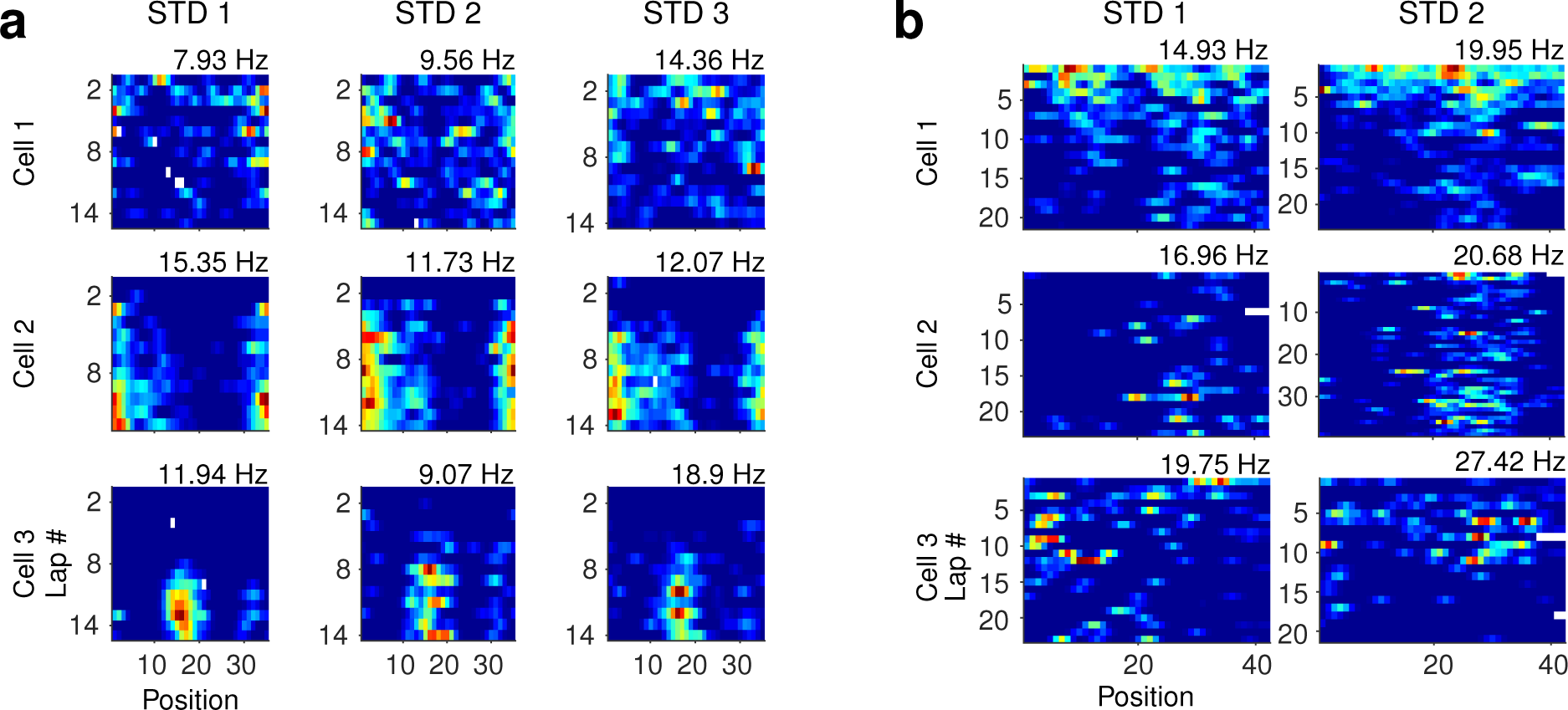
Representation of temporal information about trial identity through rate remapping of spatial firing fields by LEC neurons. (**a**) Three example cells. Each row shows the lap-wise spatial rate maps for an LEC neuron in three STD sessions in the double rotation task. Cell 1 decreases its firing over the laps of the session, whereas cells 2 and 3 increase their firing from near silence on lap 1 to robust firing at specific locations in later laps. (**b**) Same as panel a for three cells in the circular track task.

**Figure 5.**
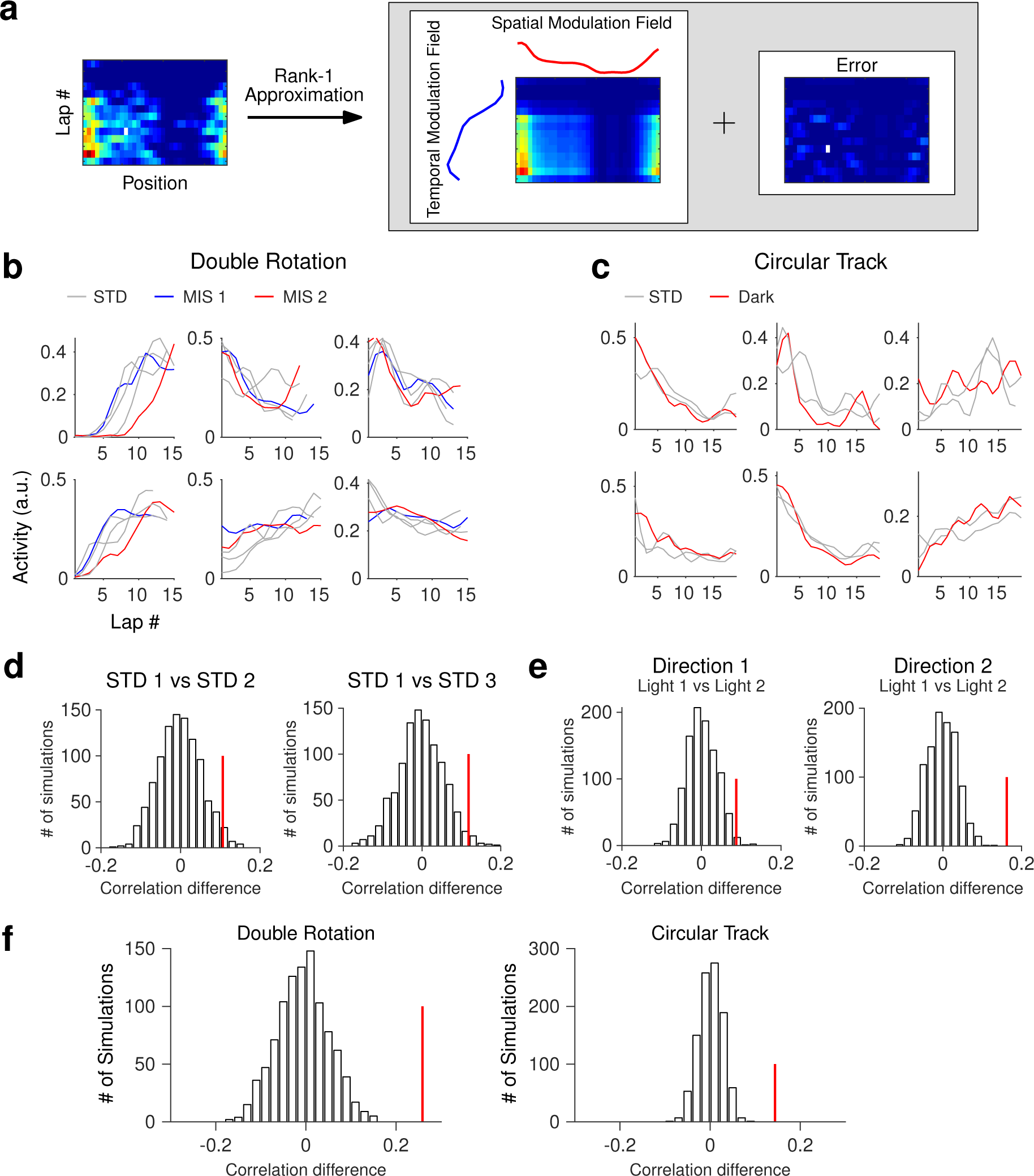
The temporal signal carried by LEC is robust to spatial manipulations. (**a**) Schematics for decomposition of the lap-wise spatial rate maps into spatial and temporal modulation fields. (**b**) The temporal modulation fields of six example cells across sessions in the double rotation task. Each cell shows similar temporal modulation in all sessions. (**c**) Same as panel b for six neurons in the circular track task. (**d**) The observed correlation (red line) and null distribution of correlations between the temporal modulation fields in two sessions in the double rotation task. Left: STD1 vs. STD2; right: STD1 vs. STD3. (**e**) Same as panel d for data in the circular track task. Left: Light 1 (STD1) vs. Light 2 (STD2) in direction 1; Light 1 (STD1) vs. Light 2 (STD2) in direction 2. (**f**) Same as panel d for correlations between temporal modulation fields in STD sessions and manipulated sessions in the double rotation task (left) and the circular track task (right).

We next investigated if the temporal signal about trial progression was robust to alterations of the spatial environment. We compared the temporal modulation fields of STD and MIS sessions in the double rotation task and the temporal modulation fields of STD and dark sessions in the circular track task. For some cells, the temporal modulation fields in STD sessions and MIS sessions were similar (Fig. 5b and 5c). The mean correlations between STD sessions and manipulated sessions were significantly larger than the null distribution created by permuting the cell identities in the second session (Fig. 5f; *p* < 0.001 for both the double rotation task and the circular track task). The temporal modulation fields were also significantly correlated between the two movement directions within a session for the circular track task and the linear track task (Fig. S6).

### Comparison of spatial and temporal representation across the hippocampal entorhinal circuit

The encoding of temporal information has been reported in the hippocampus and related regions ^1,3,42,43^, and rate remapping has also been found in both CA1 ^44,45^ and CA3 ^46^. Here, we extended our analyses to recordings from CA1, CA3, and MEC while the subjects performed the double rotation task. We computed the spatial and temporal modulation fields of neurons in the four brain regions (Fig. 6a and 6b). There were significant differences across brain regions in the spatial information scores (from the spatial modulation fields) and the temporal information scores (from the temporal modulation fields) in the first STD session (Fig. 6c and 6d; Kruskal-Wallis test; spatial information: χ^2^ = 342.86, *p* < 0.001; temporal information: χ^2^ = 15.7, *p* = 0.0013). The four brain regions, ranked in the order of spatial selectivity, were CA1, CA3, MEC, and LEC (median spatial information score, LEC: 0.36, MEC: 0.57, CA3: 1.49, CA1: 1.75; post-hoc Wilcoxon rank-sum tests, *p* < 0.001 for all tests, corrected for family-wise error rate with the Holm-Bonferroni method). (Note that the difference between CA1 and CA3 would be affected by the recording location along the transverse axis of these regions, which was not controlled for in these experiments; ^47–49)^. In contrast, LEC showed stronger temporal information coding than the other three brain regions (median temporal information score, LEC: 0.14, MEC: 0.099, CA3: 0.092, CA1: 0.10; post-hoc Wilcoxon rank-sum tests, LEC vs. MEC: *Z* = 2.65, *p* = 0.008, LEC vs. CA1: *Z* = 2.42, *p* = 0.015, LEC vs. CA3: *Z* = 3.25, *p* = 0.001; with Holm-Bonferroni corrections), and CA1 was stronger than CA3 (Wilcoxon rank-sum test, *Z* = −2.71, *p* = 0.007), consistent with earlier reports ^24,50,51^. We also compared the correlation coefficients between temporal modulation fields of standard sessions across these regions. There were significant differences between these regions (Fig. 6e; Kruskal-Wallis test, χ^2^ = 15.53, *p* = 0.0014; median correlation coefficient, LEC: 0.13, MEC: 0.02, CA3: 0.08, CA1: 0.02; post-hoc Wilcoxon rank-sum tests, LEC vs. MEC: *p* = 0.004, LEC vs. CA1: *p* = 0.0071, CA1 vs. CA3: *p* = 0.0037, corrected for family-wise error rate with the Holm-Bonferroni method).

**Figure 6.**
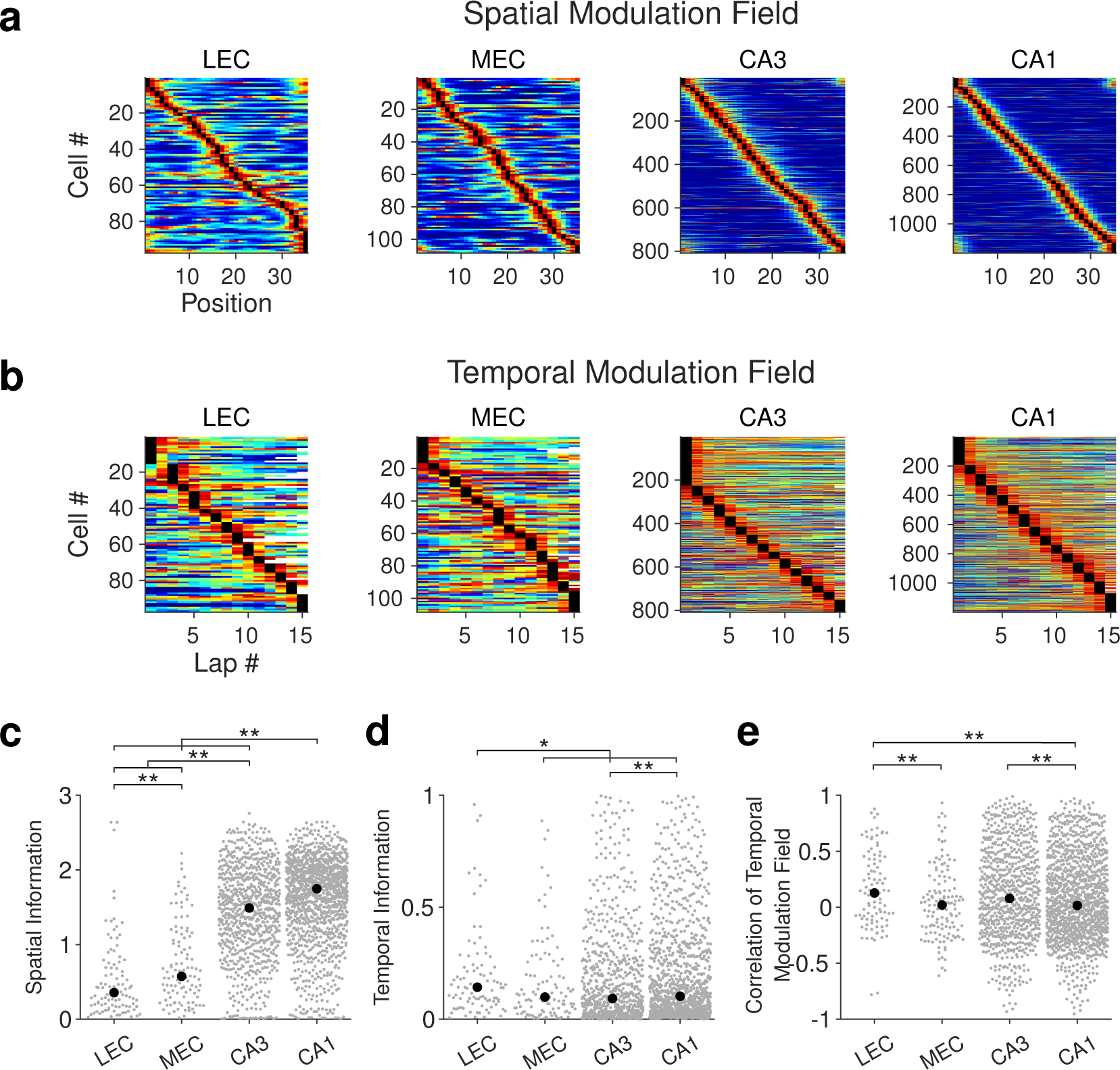
Spatial and temporal coding properties in the entorhinal-hippocampal regions. (**a**) Sorted spatial modulation fields for LEC, MEC, CA3, and CA1. (**b**) Sorted temporal modulation fields for the four brain regions. (**c**) The distributions of spatial information scores across regions. Gray dots, data for each neuron; black dots, the median value. (**d**) The distributions of temporal information scores across regions. (**e**) The distributions of Pearson’s correlation coefficients between standard sessions across regions. *, *p* < 0.05; **, *p* < 0.01.

## Discussion

In this study, we investigated the representation of spatial information and temporal information about trial progression in LEC. We found that a portion of LEC cells were selective for spatial location on linear or circular tracks. Although the spatial selectivity of LEC was generally weaker than that of MEC and other hippocampal regions ^6,15,27^, a proportion of cells showed spatial precision and reliability similar to neurons in the hippocampal regions and the MEC. The degree of spatial tuning in LEC on these 1-dimensional tracks was not previously reported in LEC cells during 2-dimensional, open-field foraging tasks in the absence of objects ^15,16^. Moreover, in 1-dimensional shuttling tasks, many LEC neurons exhibited directional and distance selectivity in their spatial firing. Within a session, the firing rate of the spatial firing fields often showed temporal modulation. This coding of trial progression in LEC was robust to manipulations of the spatial environment, such as cue rotations and the absence of visual inputs. Finally, this temporal coding through spatial rate remapping was stronger in LEC compared to MEC, CA1, and CA3.

### Spatial coding in LEC

Previous studies have examined the nature of spatial representation in LEC ^15,16,20–22,27^. In 2D empty arenas, the allocentric spatial tuning was weak ^15,16^, but in the presence of objects, a subset of LEC neurons showed greater selectivity both around the objects and away from the objects ^20^. A small number of LEC cells fired in locations that were previously occupied by objects ^20,21^, thereby demonstrating memory traces of the object’s previous locations ^21^. In the current study, the presence of a reliable spatial code may perhaps be attributed to the surface cues on the track in the double rotation paradigm, as well as the reward near the end of the journey in the circular and linear track shuttling tasks providing stable, local anchoring points for the LEC spatial signal. The persistence of spatial firing in the dark suggests that the spatial code in LEC could be supported by either olfactory input or other cognitive variables, or perhaps from path integration signals provided by the MEC inputs to LEC. Consistent with a previous study analyzing the same data, we have confirmed that the spatial representation in LEC rotated with local cues ^27^. This finding suggests that LEC may be involved in biasing the responses of CA3 neurons to local cues ^29,48^ and in segmenting the firing of CA1 neurons based on local surface boundaries ^52^.

A potential factor contributing to the discrepancies between our results and the reported lack of spatial coding in earlier studies in the double rotation task using the same dataset ^16,27^ could be our identification and removal of all head scanning movements (including the initiation and ending of head scans when the rat’s head was still over the track) in our treatment of positions, using tools developed since the publication of the Neunuebel et al. (2013) paper ^28^ (see Methods). Unlike our approach, Neunuebel et al. ^27^ only removed the off-track portions of head scans, which left potentially nonspatially modulated spikes fired during the starts and ends of head scans in the rate maps. Furthermore, because LEC cells fire in an egocentric frame of reference ^22^, the firing of LEC cells during head scanning may occur at different allocentric locations on the track when the rat’s head is at the preferred egocentric bearing relative to its anchor point in the environment. Scanning constitutes an overt attentive behavior, during which the animal may gather information from its external environment. LEC plays a role in processing multimodal sensory information ^53^, with a significant number of LEC cells displaying activity during head scanning events at various locations (Fig. S2a). The removal of this spatially nonselective activity from the rate maps enhanced the quality of spatial information represented in the LEC firing rate maps. This allowed us to reveal a higher degree of spatial modulation of LEC cells then previously reported (albeit still less than that observed in MEC and hippocampus).

An unresolved question from these data is whether the spatial coding on the 1-D tracks is allocentric or egocentric. Wang et al. ^22^ demonstrated that in 2-D open fields, LEC cells represent the animal’s location in an egocentric frame of reference, that is, as an egocentric bearing and distance to a specific point in the environment (such as the center or nearest boundary, an object, or a goal location). In contrast, MEC cells represented location in the classic allocentric frame of reference associated with spatial representations of place cells in the hippocampus. Because of the behavioral constraints imposed by the rat’s trajectories on linear and circular tracks, we do not have adequate sampling to determine whether the spatial firing of LEC in the present paper indicates an egocentric or allocentric representation. We speculate that, consistent with the results in open fields, the spatial firing is egocentric (e.g., the cell fires at a particular bearing relative to the goal location or relative to a salient external landmark). The geometric constraints on the rat’s behavior limit the sampling of specific egocentric bearings at specific allocentric locations, perhaps allowing more reliable firing of LEC neurons at specific locations compared to the more unconstrained trajectories and combinations of egocentric bearings and allocentric locations available in open fields. Resolving this question will require further experiments with proper controls to tease apart allocentric from egocentric coding in such 1-D tasks.

### Direction selectivity in LEC

The directional movement on one-dimensional tracks generates a strong context signal that influences the spatial representation within the hippocampal-entorhinal circuit ^31–36,54,55^. To our knowledge, our results constitute the first evidence of LEC neurons displaying a directional preference during shuttling tasks on a one-dimensional apparatus. This finding contributes to the existing evidence that LEC neurons might carry context-related information ^19,56,57^. Given the rich bidirectional connections of LEC with the hippocampus and MEC ^58,59^, it is conceivable that the directional maps created by hippocampal or MEC neurons could contribute to the observed direction selectivity in LEC. Nevertheless, certain features may be unique to LEC cells: (1) the directional preference of some LEC neurons spanned a large proportion of the track; (2) the directional preferences showed a concentration around the goal, possibly indicating leave- or approach-related egocentric preferences ^22,23^; (3) the selectivity for movement direction might partly result from the distance coding exhibited by some of the LEC cells. These features might aid in maintaining and stabilizing the differential representation of contexts in the hippocampal navigation circuits, and contribute to the manifestation of distance coding properties in the downstream hippocampus under specific conditions ^60^. In conclusion, our results suggest that spatial representations for travel directions are present across LEC, MEC, and hippocampal regions.

### Temporal coding through spatial rate remapping in LEC

The hippocampus is essential for processing nonspatial information integral to episodic memory ^45,61^. A proposed coding scheme involves the combination of spatial and nonspatial coding via rate modulation of the spatial firing profile, i.e., rate remapping ^45,46,62^. LEC has been shown to be involved in rate remapping of CA3 cells ^56^. To our knowledge, our study provides the first evidence that LEC cells could signal temporal information about trial progression using the same feature-in-place type of rate remapping mechanism. Note that the differences in temporal coding between LEC and other brain regions were smaller than those reported by Tsao et al. ^24^. One possibility is that the double rotation task involved more stereotypical behaviors than the open field foraging task in two-dimensional boxes used in that study. The structured behavior could diminish temporal information about trials or sessions ^24^, thus obscuring the distinctions among brain areas. In tasks involving the repeated traversal of one-dimensional tracks, we demonstrated LEC carried more minute-scale temporal information than in MEC, CA1, and CA3, which supports the view that LEC plays an essential role in coding time information ^24,25^. Furthermore, the reset of network pattern to a fixed initial state at the beginning of a session and the consistent temporal modulation of the spatial firing of LEC neurons across different sessions, even after considerable contextual changes, can help to suppress arbitrary drift across sessions in other cortical areas ^63^. Consistent with this proposal, inactivating LEC led to decreased representational consistency in the medial prefrontal cortex for stimuli ^64^. Our results are in line with previous reports that CA1 neurons could use rate remapping to represent temporal information ^44,51,65–67^. Together, our results suggest that, as one of the major inputs to the hippocampus, LEC may contribute to the representation of time in the hippocampus ^24,25^.

In conclusion, our data have demonstrated reliable spatial processing in LEC neurons, and LEC neurons represent temporal information about trial progression through spatial rate remapping. The temporal modulation of place fields, together with time cells, ramping cells, representational drift, etc., provide a hierarchical framework to code temporal information. The integration of space and time processing in LEC and other areas suggests a shared organizational feature across the MTL memory system ^2,3,43,68^.

## Acknowledgments

STI2030-Major Projects 2022ZD0205000

NIH Grant R01 MH094146

NIH Grant R01 NS039456

Shenzhen Key Laboratory of Precision Diagnosis and Treatment of Depression (ZDSYS20220606100606014)

Chinese Academy of Sciences Pioneer Hundred Talents Program

Guangdong Basic and Applied Basic Research Foundation (2021A1515010809)

National Natural Science Foundation of China (32171043)

CAS Key Laboratory of Brain Connectome and Manipulation (2019DP173024)

Guangdong Provincial Key Laboratory of Brain Connectome and Behavior (2017B030301017)

## Author contributions

J.J.K. and C.W. conceived the study. J.J.K. supervised the experiments. C.W., H.L., and G.R. performed recording experiments. C.W. performed data analyses. C.W. and J.J.K. wrote the first draft of the manuscript. All authors commented on the manuscript.

## Materials & Correspondence

Correspondence and material requests should be addressed to J.J.K. or C.W.

## Methods

### Subjects and surgical procedures

#### The circular and linear track paradigm

Five male, 5-6 months old, adult Long-Evans rats obtained from Envigo performed the circular track and linear track tasks. The animals were housed individually on a 12:12 h reversed light-dark cycle. Experiments were carried out in the dark phase of the cycle. Animal care, surgical procedures, and euthanasia complied with NIH guidelines and were approved by the Institutional Animal Care and Use Committee at Johns Hopkins University. Detailed surgical procedures were reported elsewhere ^22^. Briefly, a hyperdrive targeting LEC in the right hemisphere was implanted (7.55-7.6 mm posterior to bregma, 3.0 mm lateral to the midline) with the tetrode bundles angled at 25° mediolaterally. Rats were food-restricted until their body weight reached 80-90% of their free-feeding weights. The tetrodes were lowered slowly toward LEC over the course of 1-2 weeks.

#### Double rotation paradigm

Neurons from 60 rats in the double rotation task (3 LEC rats, 6 MEC rats, 32 CA1 rats, and 29 CA3 rats; in some rats two areas were recorded simultaneously) were analyzed using previously published data sets. Details of the surgical procedures conducted on these rats can be found in previous papers ^16,27,29,30,48^.

#### Electrophysiology and recording

Tetrodes were made from 12 μm or 17 μm nichrome or platinum-iridium wire (California Fine Wire, Grover Beach, CA, USA or Kanthal, Palm Coast, FL). The electrode tips were electroplated with gold to 200-500 kOhm with 0.2 µA current or 120 kOhm with 0.075 µA plating current. Platinum iridium wires were not plated and had an impedance of ∼700 kOhms. During recordings, the electrophysiological signal was either processed by a 64-channel wireless transmitter system (Triangle Biosystems International, Durham, NC) or a unity-gain preamplifier headstage (Neuralynx, Bozeman, MT). The data were then collected with Cheetah Data Acquisition System (Neuralynx, Bozeman, MT). For single unit recordings, the signal was band-pass filtered (600 Hz to 6 kHz) and thresholded (∼70 µV) to produce the waveforms of the spikes. We used red and green LEDs or arrays of infrared LEDs on the head of the subjects to track the positions of the animals, and a color or infrared CCD camera with a 30 Hz sampling rate was used to capture the color or infrared signals, respectively. The behavioral trajectories of the rats were smoothed with a 5-frame boxcar filter (150 ms) and speed filtered (3 cm/s).

#### Histological processing procedures

The rats were perfused transcardially with 4% formalin. The brains were extracted and submerged in a 30% sucrose formalin solution. Brain tissue was then cryosectioned at 40 µm thickness and mounted onto glass slides. Standard Nissl staining was performed to identify the tetrode tracks. Free-D software ^69^ was used to register tetrode tracks to the tetrode bundle configuration of the drive. Recording locations of each session were determined based on the amount of tetrode turning each day and histological reconstructions of electrode tracks ^15^.

The demarcations of LEC followed conventions in previous studies (Fig. S1)^70–72^. Briefly, LEC was distinguished by the large and darkly stained neurons that formed discontinuous islands in layer II, and the presence of a cell-sparse lamina dissecans between layer III and layer V.

For details on the identification of MEC, please refer to a previous study ^27^.

### Behavioral Tasks

#### The circular track and linear track task

Animals were trained to run back and forth between food wells for food pellets (BioServ) on a circular track (diameter 97 cm, width 10 cm). The food wells were separated by a 15 cm tall black barrier with 0.4 cm sidewalls; thus, the animals needed to run almost 360° in each direction to obtain the next food reward. The linear track task was similar to the circular track task, except the apparatus was a 5-foot long, 8-cm wide linear track. Each session consisted of 7 to 25 laps. Part of the data has been reported in a previous paper (supplementary Fig. 9 of ^24^).

#### The double rotation task

The data collected in the double rotation task have been reported for other purposes ^27–30,48^. Briefly, the animals were trained to run on a circular track with local texture cues in the clockwise direction (outer diameter: 76 cm, inner diameter: 56 cm). The subjects foraged for food scattered at arbitrary locations on the track (∼2 rewards/lap) while moving clockwise for about 15 laps. The apparatus was surrounded by a black curtain with salient cues. The experiment lasted for four days and was conducted during the dark portion of the light/dark cycle. There were two types of sessions: standard sessions (STD) and mismatch sessions (MIS). In STD sessions, the relationships between local and global cues remained fixed; in MIS sessions, local and global cues were rotated by an equal amount in opposite directions, thus generating mismatch angles of 45°, 90°, 135°, and 180° (only 135° and 180° were used here for analysis of spatial coding properties, as we concentrated on the two most extreme manipulations). Each session consisted of about 15 laps. Rats either ran five sessions (STD, MIS, STD, MIS, STD) or six sessions (STD, STD, MIS, STD, MIS, STD). In the case of the six-session configuration, the data in the first STD were considered as the baseline, and not analyzed in the current paper.

### Unit isolation

Single units were isolated manually with custom-written spike-sorting software (Winclust, developed by J. Knierim). The peak amplitude and energy of the waveforms were used to isolate cells. The quality of each unit was rated with a score ranging from 1 (very good) to 5 (poor). The cluster isolation quality was assigned completely independent of any behavioral correlates of the cells. Units rated as 4 or 5 were excluded from the analysis.

### Data analysis

All analyses were performed with Matlab. All statistical tests were two-sided.

### Linearization of the trajectories and detection of head scanning events

In the linear track paradigm, the long edge of the apparatus was aligned with the vertical dimension in the camera’s field of view. Therefore, we linearized the position of animals’ trajectories with only the vertical data. In the circular track paradigm and double rotation paradigm, we transformed the data to a polar coordinate with the center of the track as the origin.

In the double rotation paradigm, head scanning events were detected following a previous study^28^. Briefly, head scanning events were defined as lateral head movements off the track with a minimum duration of 0.4 s and a minimum radial extent of 2.5 cm. These behavioral epochs and spikes associated with them were removed from the analyses. See Monaco et al. ^28^ for a detailed description of these procedures.

### Spatial firing rate maps

The spatial bin was set to 6 cm, and a Gaussian kernel (sigma: 1 bin) was used to smooth the final rate map. By contrast, in Neunuebel et al., ^27^, the bin size and the sigma of the Gaussian kernel were 0.58 cm and 3.1 cm, respectively. The spatial information score was calculated following a previous study ^73^.

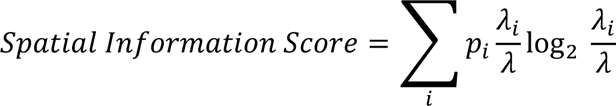

where *p*_i_ was normalized occupancy in the *i*-th spatial bin, *λ*_i_ was the spatial response (firing rate) in the *i*-th bin, and *λ* was the average response.

### Spatial and temporal modulation fields

To estimate the spatial and temporal firing properties of neurons, we applied singular value decomposition (SVD) to the lap-wise spatial rate map matrix formed by stacking spatial rate maps from each lap. Any lap that had less than 70% coverage of the track was removed as in some laps the animals turned back before they reached the target. The remaining unvisited spatial bins were set to 0 before performing the decomposition. The resulting Euclidean norm (vector length) of both the spatial and temporal modulation fields was 1. The temporal and spatial modulation fields were smoothed with a Gaussian kernel (standard deviation: 1 bin). The information score was calculated from the spatial and temporal modulation fields in the same way as the section above.

### Correlation matrix

The population correlation matrix was constructed following previous studies ^48^ for both spatial and temporal rate maps (or modulation fields). For each cell, we first normalized the rate maps by the peak of the two spatial rate maps to be compared. The normalized firing from all cells for a bin was used to create a population vector. Population vectors for every bin were correlated to those in the second session using Pearson product-moment correlation. The resulting matrix was termed the population correlation matrix. In this study, the correlation matrix method was only used as a visualization tool.

### Permutation test for consistency of temporal and spatial modulation across sessions

To characterize the similarity of spatial or temporal firing patterns between two sessions, for each cell, we calculated Pearson correlation coefficients between a pair of firing maps or modulation fields and obtained the mean correlation of the population. Then, the cell identities in the second session were randomly permuted, and the average correlation was recomputed for these random pairs. This operation was repeated 1000 times to build a null distribution of correlation values. The observed average correlation coefficient was then compared to a null distribution with a significance level of 0.05.

### Direction selectivity analysis

To determine if cells show significant direction selectivity on regions in the one-dimensional track, we adopted the nonparametric permutation method of Fujisawa and colleagues ^37^ to set the threshold for significant differences between firing in the two movement directions. The lap-wise spatial rate map was obtained separately for two movement directions. We obtained averaged rate maps across these laps to get the session-wise mean firing and their differences at each spatial bin. Then the labels of the movement direction were permuted, and the differences in firing rate for the resampled groups were calculated. These operations were repeated 500 times. For each spatial location, both the pointwise and the global acceptance bands were obtained. The pointwise acceptance bands were defined as the 97.5 percentile and 2.5 percentile of the resampled null distribution. To maintain a familywise error rate of 5% for detecting the presence of directional firing at a location, we calculated a global acceptance band as follows. We increased the pointwise acceptance bands defined from the original 500 resampling statistics in steps of 0.1 percentile. We repeated the permutation operation 500 times for each step, until the number of cases that crossed the upper and lower acceptance bands at any spatial bin was fewer than 25 times (5% of the total number of simulations). The resulting percentiles defined the global acceptance band. If the difference in firing between the two directions crossed outside the global acceptance bands, then the cell was considered to be directionally selective at this location. We then used the pointwise acceptance bands to define the size of the direction-selective region. For each global crossing, we measured the extent of the difference in firing between the directions that remained outside the bands of the pointwise acceptance bands.

The direction selectivity index was defined as:

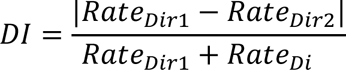

where *Rate*_Dir1_ and *Rate*_Di_ represent the firing rates at a particular position for two movement directions. The possible range of *DI* is 0 (no direction selectivity) to 1 (cell only fires for one direction, i.e., strongest possible direction selectivity).

### Test for distribution of direction-selective regions

For the circular track data, we converted the peak of the direction-selective region to polar coordinates, and the Rayleigh test was used to test for the non-uniformity of the resulting circular data. For the linear track data, we performed a Monte Carlo test. We divided the track into two regions: the center region and the end region. The end region was the portion of the track close to either track end (a quarter of the whole track). The center region occupied half of the track in the center. We calculated the proportion of the data that belongs to the end regions and compared the proportion both from real data and randomly generated numbers (1000 times, null distribution). The data was considered to have a significant preference to concentrate close to the end if the observed proportion was significantly larger than the 95^th^ percentile of the null distribution.

### Test for distance coding properties of LEC neurons in shuttling tasks

We averaged the values along the minor diagonal of the population correlation matrix created with the spatial rate maps for the two movement directions. This mean correlation was compared to the null distribution created by permuting the cell identity in the opposite direction 1,000 times.

To test whether there was an equal amount of distance tuning in the starting and ending position of the journey, the starting position was defined as the first 4 spatial bins and the ending position was the last 4 spatial bins. We calculated the observed difference of correlation between the two regions and compared that value with a simulated null distribution obtained by 1,000 permutations of the direction labels. The significance level was 0.05.

### Data availability

The data used in this study are available from the corresponding author upon reasonable request.

---

**Supplementary Figure 1.**
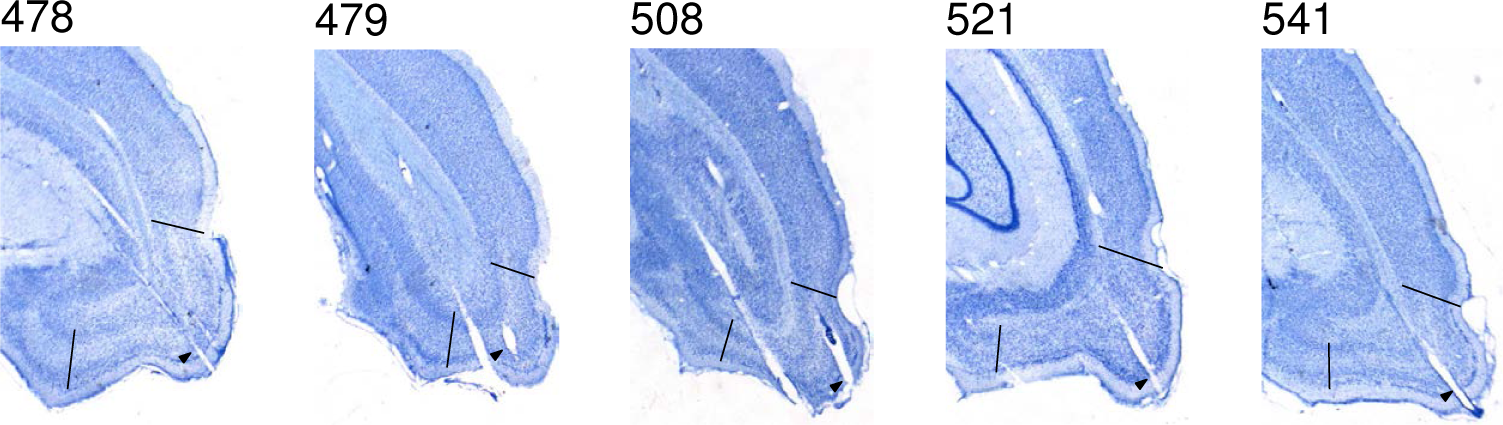
Example electrode track locations for LEC recording experiments. The black lines are the medial and lateral boundaries of the LEC. The arrowheads denote example tetrode tracks.

**Supplementary Figure 2.**
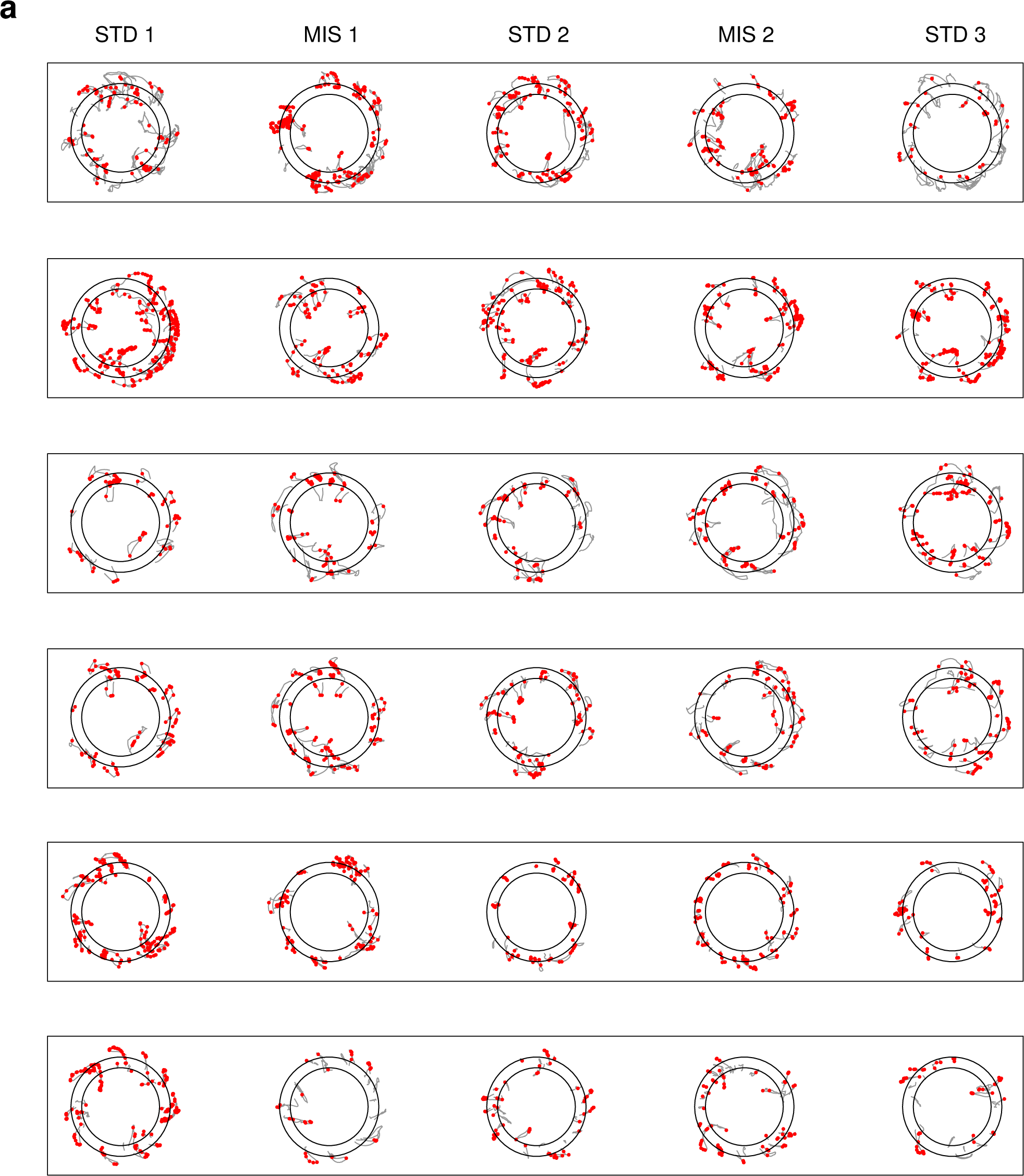
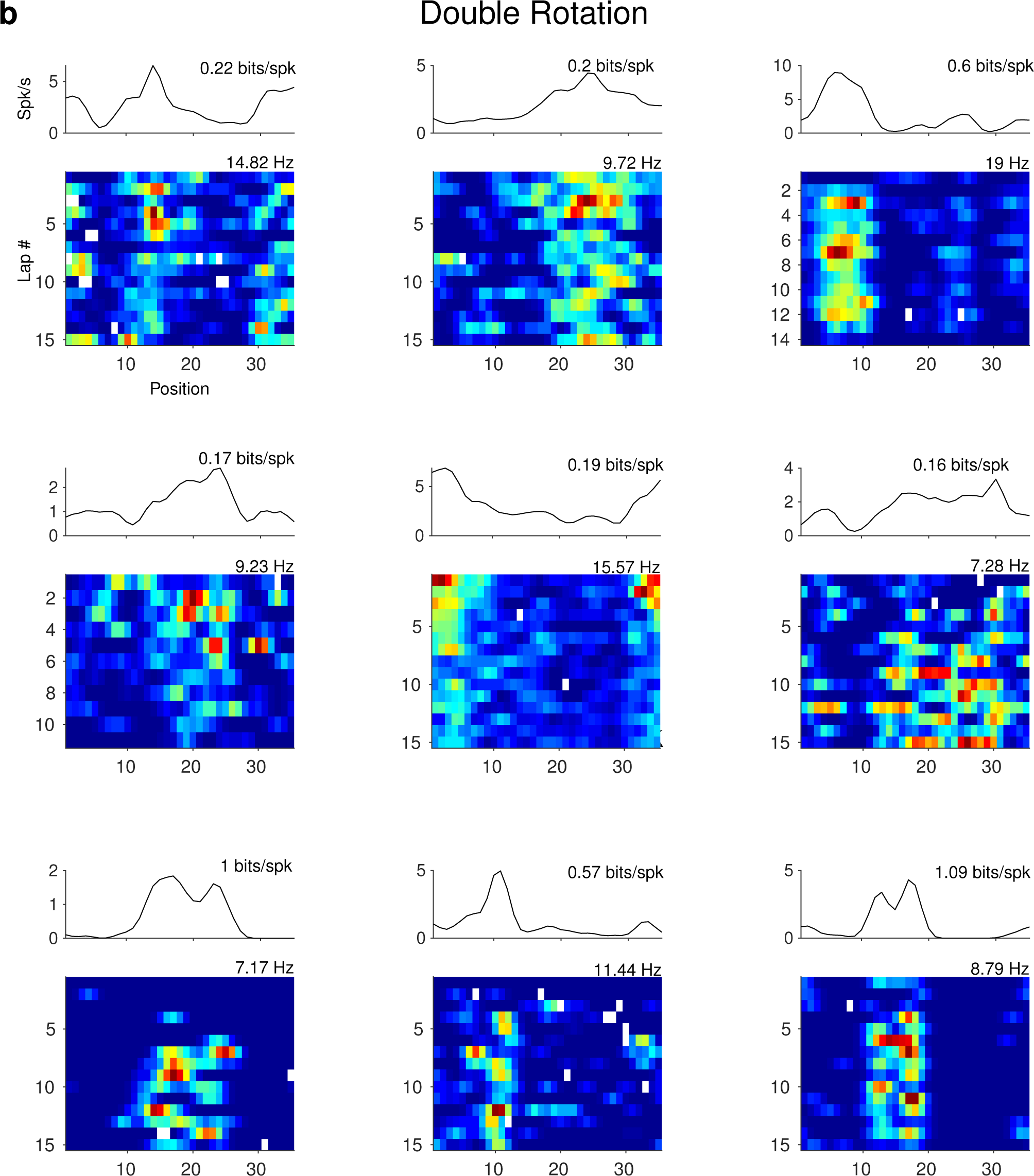
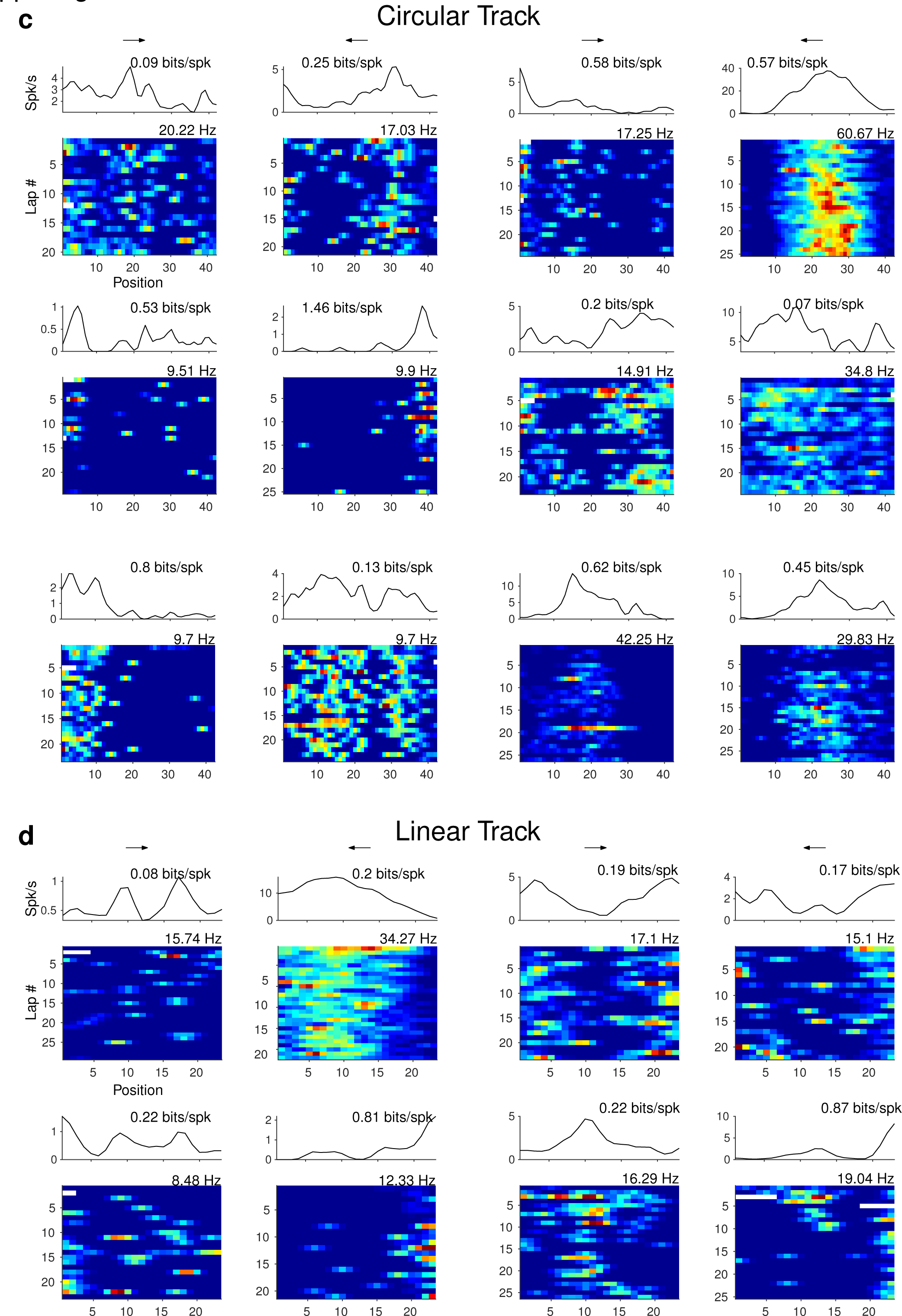
Example spatial trajectories and spatial rate maps. (**a**) Example head-scanning events and head scanning-related firing of 6 LEC neurons in the double rotation task. Black circles: the inner and outer contours of the circular track. Gray lines: the trajectories of the subjects. Red dots: The locations of the subject when firing occurred. In the current study, the complete extent of each scanning event and the associated firing were identified and removed, which may explain why we uncovered greater spatial selectivity in some LEC neurons compared to previous reports with this data set. (**b**) Example spatial rate maps of LEC neurons in the double rotation task (format is the same as Figure 1d). (**c**) The circular track task (format is the same as Figure 1e). (**d**) The linear track task (same as panel c).

**Supplementary Figure 3.**
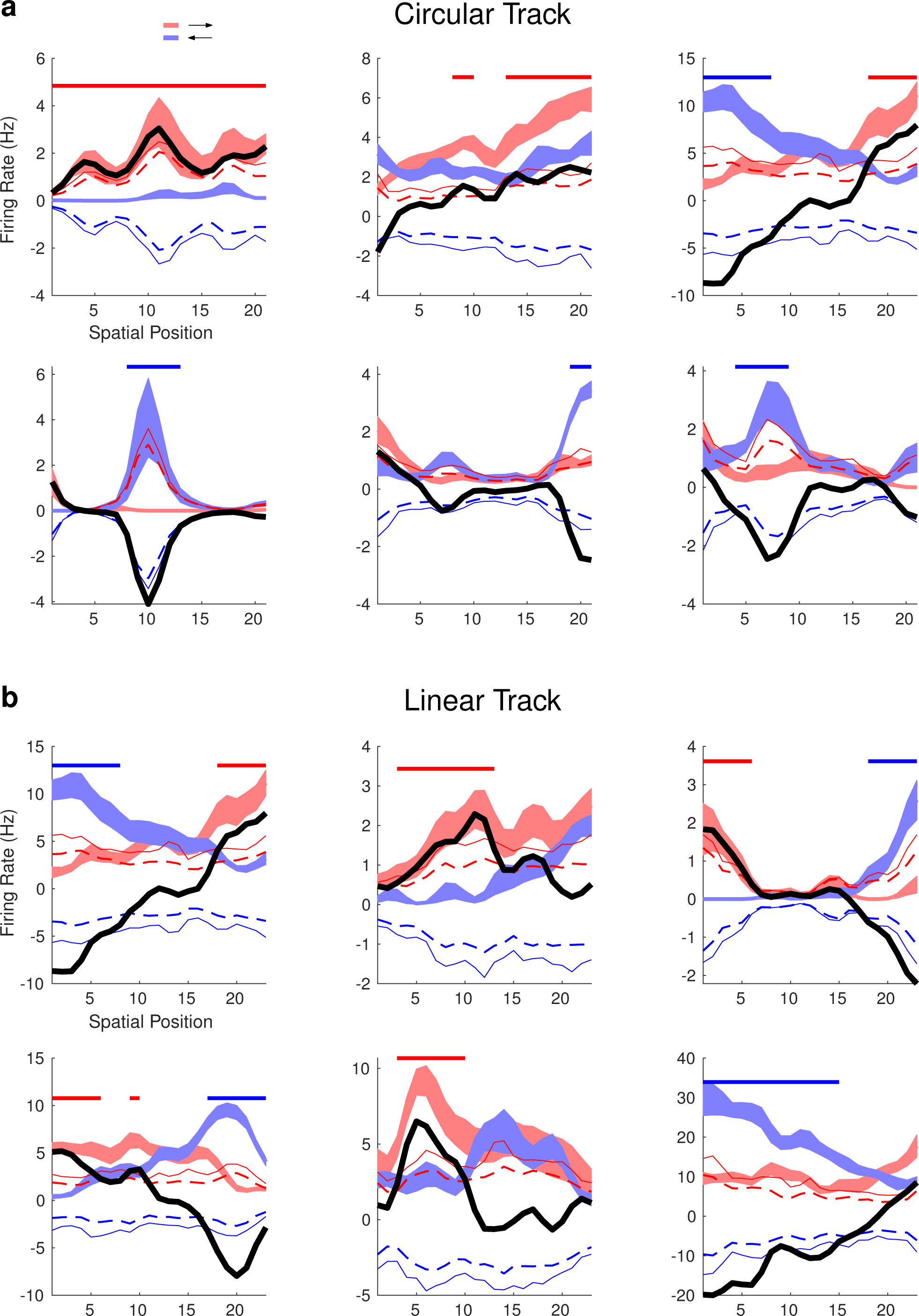
Examples of 12 LEC neurons that show direction selectivity in the circular track task (**a**) or the linear track task (**b**). The format is the same as in Figure 3e. The top left cell showed significantly greater firing for the red direction across the whole region. The left-most cell in the second row shows strong direction selectivity in the middle of the track. Distance coding was present in the neurons that preferred different directions at the two track ends.

**Supplementary Figure 4.**
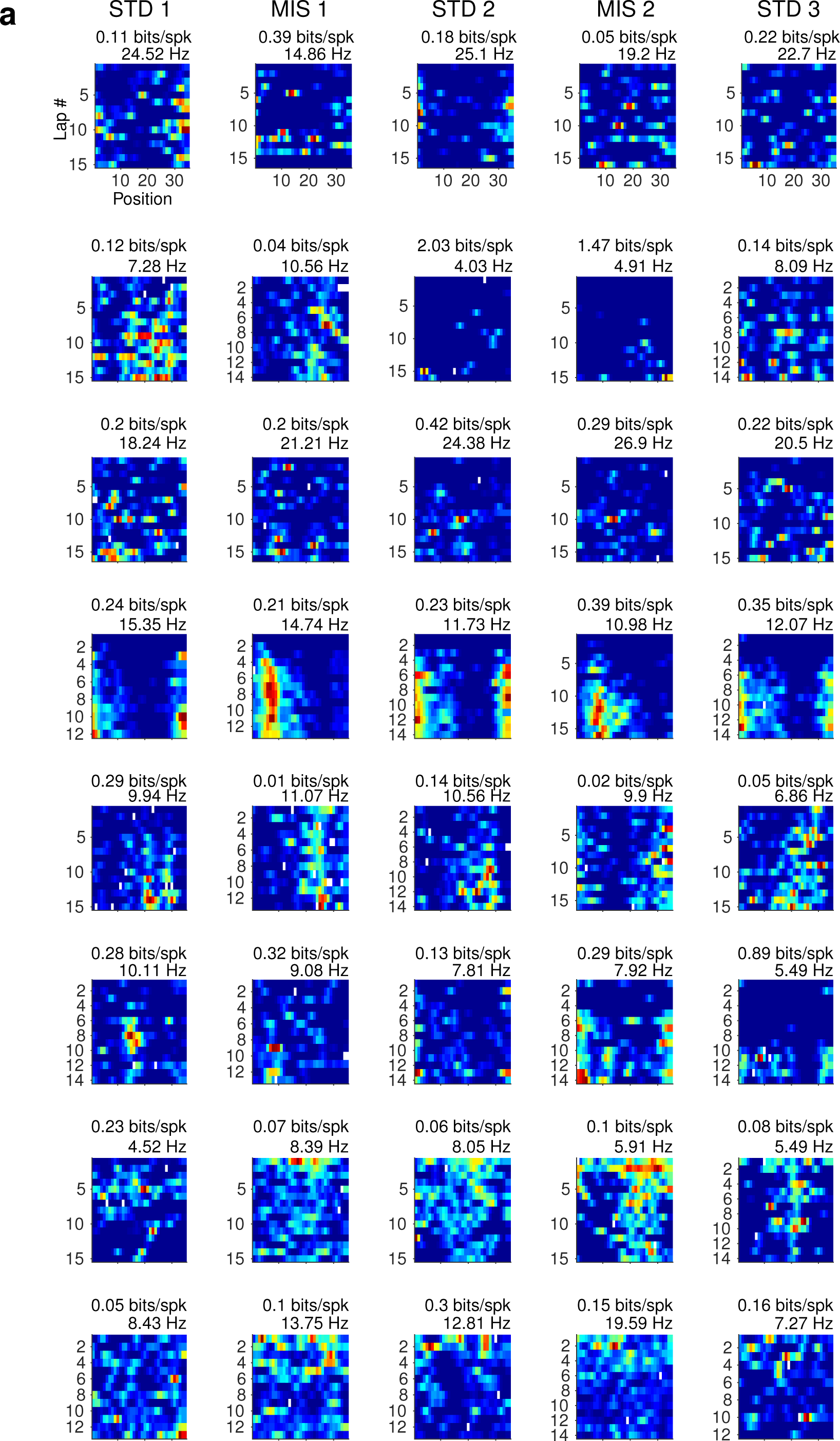
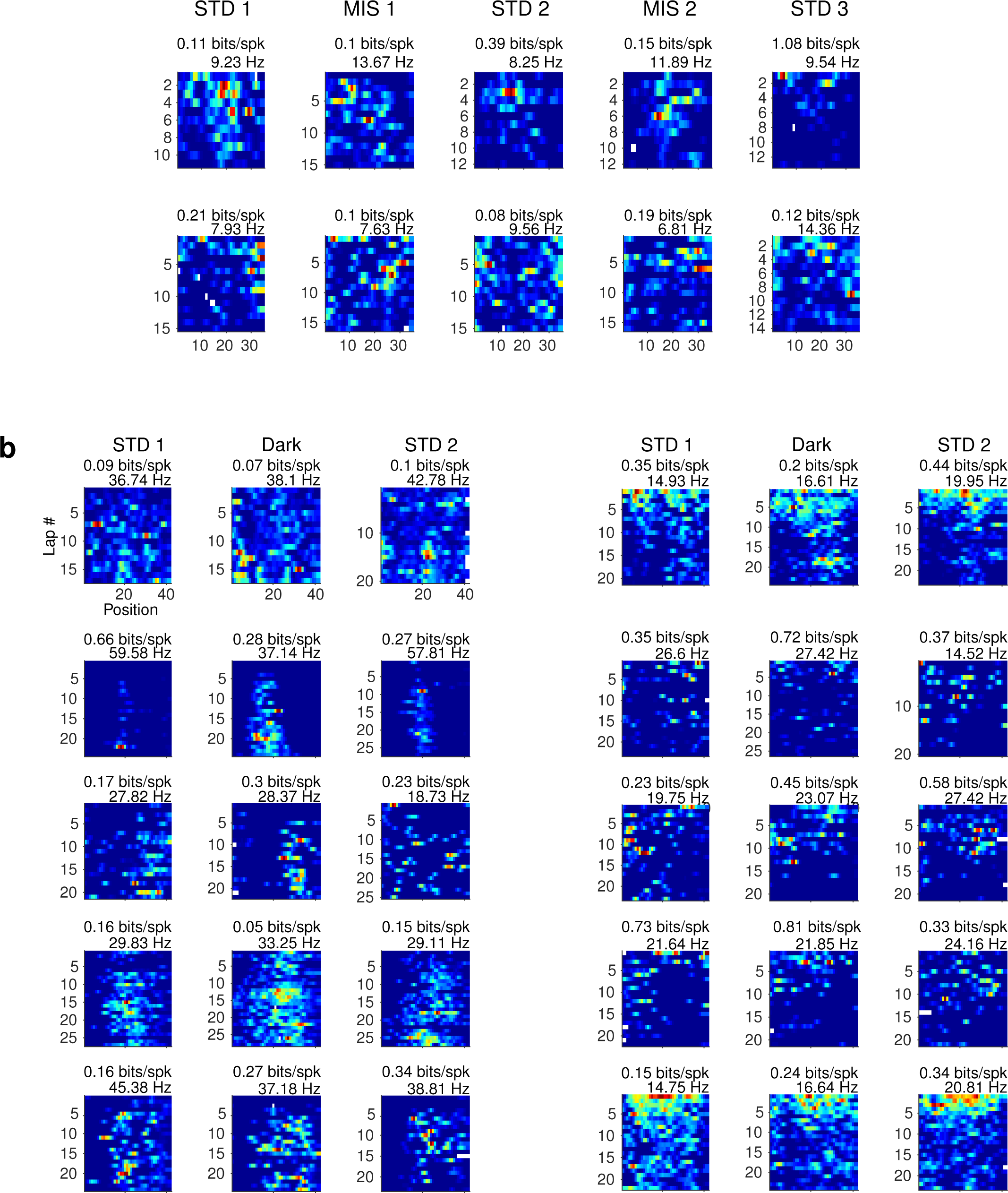
Examples of LEC cells with selectivity for trial progression. (**a**) Ten example cells in the double rotation task. Each row is a neuron, each panel is the lap-wise spatial rate map in a session. The temporal information score and peak firing rate in the rate map are shown at the top. The firing rates of the first six neurons increased gradually in the sessions, whereas the firing rate of the last four neurons decreased. (**b**) Ten example cells in the circular track task. The neurons with ascending or descending temporal profiles are shown on the left or the right side, respectively.

**Supplementary Figure 5.**
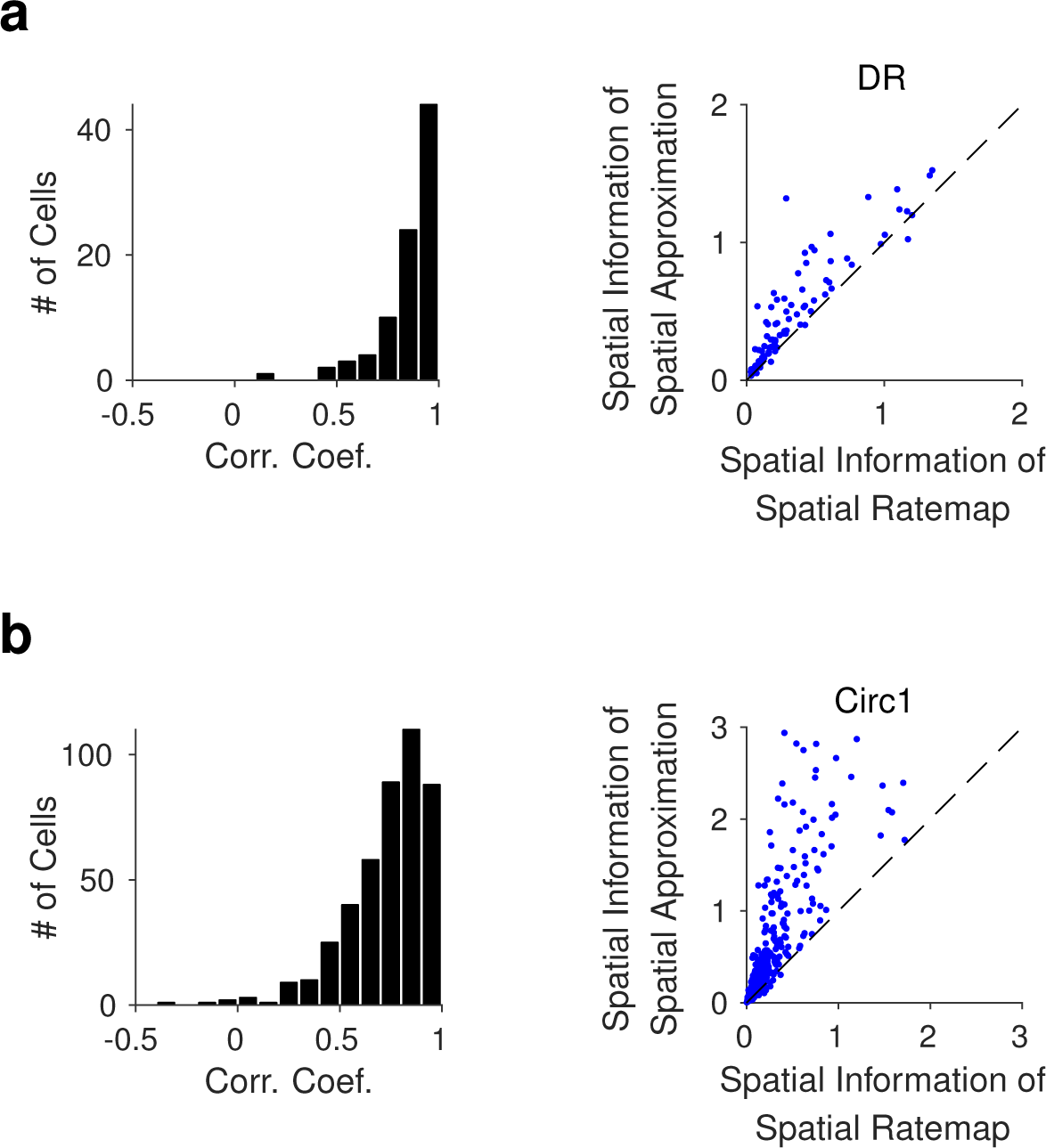
Comparison between the properties of the spatial modulation field and standard session-wide spatial rate maps (only data from the first standard session were included). The spatial information scores of the spatial modulation fields were significantly correlated with, and larger than, those from the spatial rate maps for the double rotation task (panel A; Pearson’s correlation coefficient, *r* = 0.89, *p* < 0.001; Wilcoxon signed-rank test, *Z* = −7.77, *p* < 0.001) and for the circular track task (panel B, Pearson’s correlation coefficient, *r* = 0.89, *p* < 0.001; Wilcoxon signed-rank test, *Z* = −12.43, *p* < 0.001). In some cases, the standard spatial rate map may be confounded by temporal modulations, thus resulting in inferior spatial selectivity compared to the spatial modulation fields, which is the orthogonal spatial component from the lap-wise spatial rate maps.

**Supplementary Figure 6.**
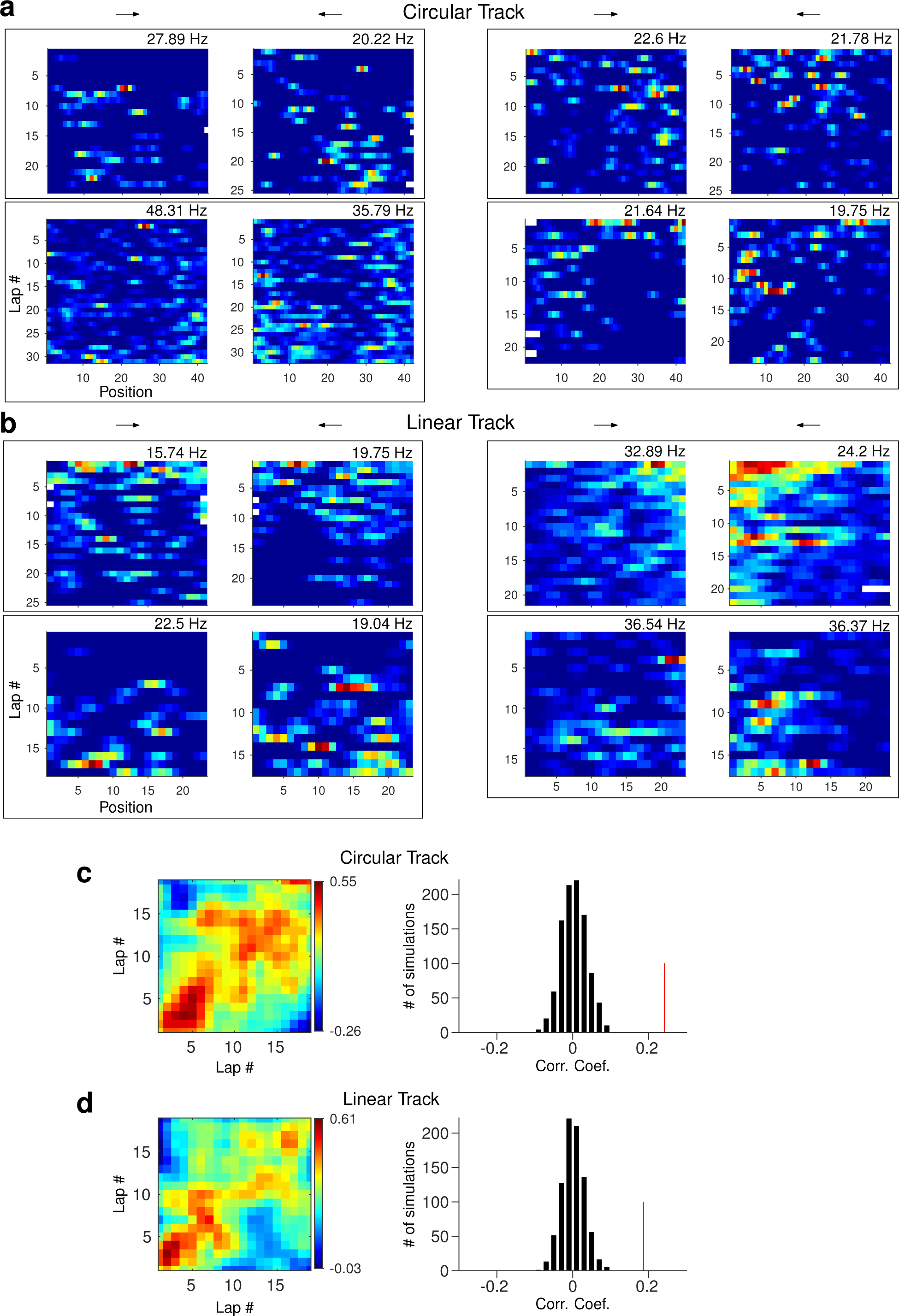
The temporal modulation fields for two directions within a session are correlated. (**a**) The lap-wise rate maps for the directions of four example LEC neurons in LEC in the circular track task. (**b**) The same format as panel a for four LEC cells in the linear track task. (**c**) Left, the population correlation matrix between the temporal modulation fields in the two movement directions; right, the observed mean correlation (the red line) between the temporal modulation fields of each cell was compared to the distribution produced by 1,000 random permutations of the data (see Methods, *p* < 0.001). (**d**) same as panel c for the linear track task (*p* < 0.001).

